# Evolution of empathetic moral evaluation

**DOI:** 10.1101/447151

**Authors:** Arunas L. Radzvilavicius, Alexander J. Stewart, Joshua B. Plotkin

## Abstract

Social norms can promote cooperation in human societies by assigning reputations to individuals based on their past actions. A good reputation indicates that an individual is worthy of help and is likely to reciprocate. A large body of research has established the norms of moral assessment that promote cooperation and maximize social welfare, assuming reputations are objective. But if there is no centralized institution to provide objective moral evaluation, then opinions about an individual’s reputation may differ across a population. Here we use evolutionary game theory to study the effects of empathy – the capacity to make moral evaluations from the perspective of another person. We find that empathetic moral evaluation tends to foster cooperation by reducing the rate of unjustified defection. The norms of moral evaluation previously considered most socially beneficial depend on high levels of empathy, whereas different norms are required to maximize social welfare in populations unwilling or incapable of empathy. We demonstrate that empathy itself can evolve through social contagion and attain evolutionary stability under most social norms. We conclude that a capacity for empathetic moral evaluation represents a key component to sustaining cooperation in human societies: cooperation requires getting into the mindset of others whose views differ from our own.

## 1 Introduction

Widespread cooperation among unrelated individuals in human societies is puzzling, given strong incentives for exploitative cheating in well-mixed populations [13]. Theories of cooperation based on kin selection, multilevel selection, and reciprocal altruism [11] provide some insight into the forces driving cooperation, but in human societies cultural forces appear to be of much greater importance [5]. One possible explanation rooted in cultural norms is that humans condition their behavior on moral reputations: the decision to cooperate will depend on the reputation of the recipient, which itself depends on the recipient’s previous actions [10]. Altruistic behavior, for instance, may improve an individual’s reputation and confer the image of a valuable society member, which attracts cooperation from others in future interactions [8].

Game theory has been used to study how reputations might facilitate cooperation in a population engaged in repeated social interactions, such as the Prisoner’s Dilemma or the Donation Game [16, 10]. In the simplest analysis an individual’s reputation is binary, either ‘good’ or ‘bad’, and the strategy of a potential donor depends on the recipient’s reputation [12] – for example, cooperate with a good recipient and defect against a bad recipient. A third-party observer then updates the reputation of the donor in response to her action towards a recipient. Reputations are governed by a set of rules, known as a *social norm*, which prescribes how to update an individual’s reputation based on the outcomes of her social interactions.

A common simplification in models of moral reputations is that all reputations are both publicly known and fully objective (e.g. [8, 15, 14, 20]). This means that all individuals know the reputations of all members of the society, and personal opinions about individual reputations do not differ. This is a reasonable assumption if there is a central institution that provides objective moral evaluation, or if individual opinions regarding personal reputations homogenize rapidly through gossip [10]. But these conditions are rare in human populations, and opinions about reputations typically differ among individuals – for instance, because observers use different moral evaluation rules, or because of divergent observation histories, or errors. In these cases a single individual may have different reputations in the eyes of distinct observers.

Moral relativity – that is, when an individual’s reputation may depend on the observer – introduces an interesting and overlooked ambiguity in how an observer should evaluate a donor interacting with a recipient. One approach is to assume that the observer can refer only to her own opinion of the recipient’s reputation, when evaluating a donor. We call this an ‘egocentric’ judgment, because the observer makes moral evaluations solely from her own perspective (Figure 1a). Alternatively, an observer can perform a moral evaluation that accounts for the recipient’s reputation in the eyes of the donor (Figure 1b). This ‘empathetic’ case requires that the observer take the perspective of another person, which assumes some capacity for recognizing the relativity of moral status. While it is well known that humans are capable of empathetic concern and perspective-taking, and that empathy may contribute to our tendency to cooperate [1, 2], the role of empathy for moral evaluation of social behavior has not been carefully studied.

Here we work to resolve the ambiguity of subjective moral judgment by introducing the concept of empathy into game-theoretic analyses of cooperation. We treat empathy E as the probability that an observer will form moral evaluations from the perspective of another person (the donor, Figure 1c). First we investigate the effects of empathy on the level of sustained cooperation under simple social norms, while players update their strategies. Next we consider evolution of empathy itself using the formalism of adaptive dynamics; and we determine conditions under which empathy will evolve and remain evolutionarily stable.

**Figure 1:**
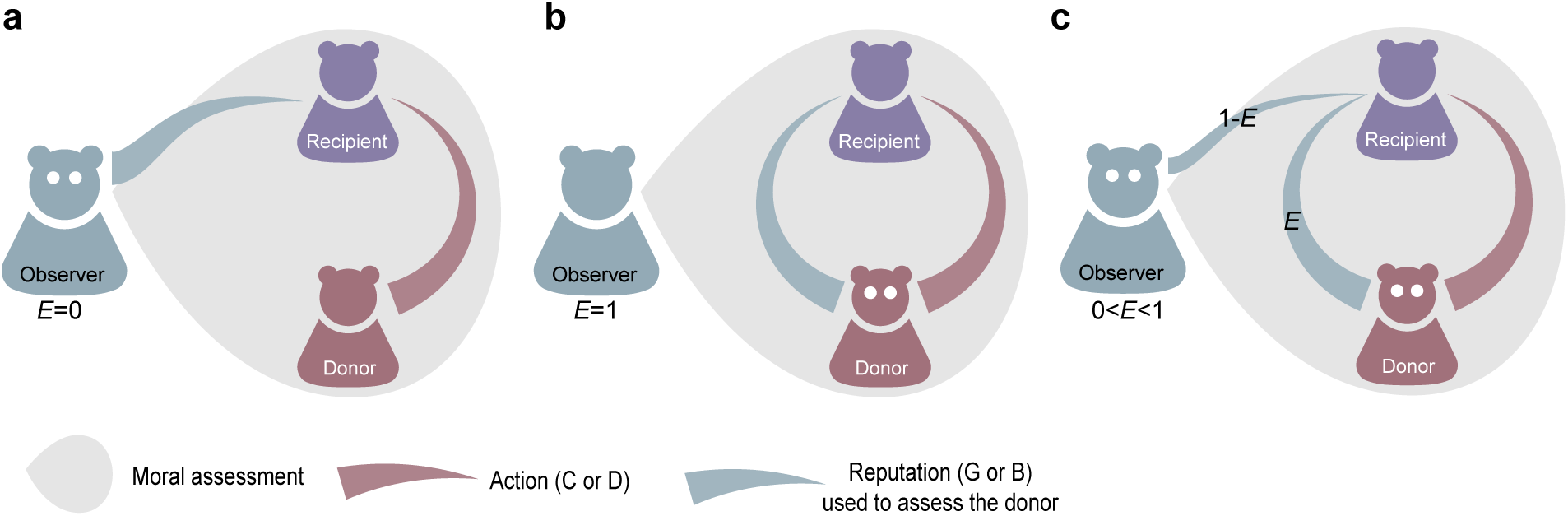
Empathetic and egocentric modes of moral assessment. An observer updates the reputation of a donor based on the donor’s action towards a recipient and the recipient’s reputation. (**a**) An egocentric observer (*E* = 0) forms a moral judgment based on the recipient’s reputation as seen from her own perspective. (**b**) An empathetic observer makes a judgment based on the recipient’s reputation in the eyes of the donor (*E* = 1). (**c**) More generally the empathy parameter *E* corresponds to the probability that observer will assess the donor using the donor’s – not the observer’s – perspective of the recipient’s reputation.

## 2 Results

### 2.1 A model of moral assessment in the donation game

We consider a population of individuals who can choose between cooperation or defection in a sequence of pairwise, one-shot donation games. In a given game the donor must choose whether or not to cooperate with the other player. If a donor cooperates she pays the cost of an altruistic act c, while the recipient receives the benefit *b* > *c*; if the donor defects she does not incur any cost, and the recipient does not receive any benefit. The donation game is therefore a special case of the prisoner’s dilemma [16] characterized by the payoff matrix 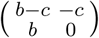.

The decision to cooperate or defect depends on the donor’s strategy *S* = [*p*,*q*], which prescribes an action conditioned on the reputation of the recipient. Here *p* and *q* denote the probability that the donor will cooperate with a ‘bad’ or a ‘good’ recipient, respectively. In our analysis we assume that both *p* and *q* are in {0,1}, and so we focus on three strategies [20]: Always Cooperate (ALLC, *S* = [1,1]); Always Defect (ALLD, *S* = [0, 0]); and Discriminate (DISC, *S* = [0,1]), which cooperates when paired with a recipient with a good reputation and defects against a recipient with a bad reputation.

Players’ reputations in the eyes of each member of the society are updated according to a social norm. In general, the update rule prescribed by a social norm can depend on the whole history of donor-recipient interactions, including the reputations of all interacting parties [19]. Complex rules of moral evaluation, however, require high cognitive ability and effort that seem unrealistic in real-world social interactions. Moreover, relatively simple “second-order” norms of moral assessment, which update a donor’s reputation based solely on the donor’s action and the recipient’s reputation, tend to outperform more complex social norms [19].

We consider second-order social norms, which can be encoded by a binary matrix *N_ij_*. The row-index *i* indicates donor’s action, *i* = 1 for defect or *i* = 2 for cooperate; and the column-index *j* indicates reputation of the recipient, *j* = 1 for bad or *j* = 2 for good. We focus on the four second-order norms that are most prominent in the literature: Stern Judging 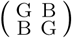, Simple Standing 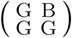, Scoring 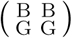, and Shunning 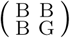. For example, under Stern Judging (SJ) or Simple Standing (SS) an observer will assign a good reputation to a donor who punishes a recipient with a bad reputation, by defection. Whereas under Shunning (SH) or Scoring (SC) an observer will assign a bad reputation to a donor who defects against any recipient, regardless of the recipient’s reputation. Following [20] we also allow for errors in strategy execution and in observation: a cooperative act is erroneously executed as defection with probability *e*_1_, while an observer erroneously assigns bad reputation instead of a good reputation, and *vice versa*, with probability *e*_2_.

The broad consensus in the literature is that Stern Judging is the most efficient norm for promoting cooperation, along with widespread adoption of the discriminator strategy. This result is robust with respect to strategy exploration rates [17], population sizes and error rates [18], and it even extends to the realm of more complex norms of third- and fourth-order [19]. Pacheco et al. [15] have additionally shown that Stern Judging is the norm most likely to evolve in a model of group-structured population, because it maximizes the collective payoff of the group.

However, prior studies of cooperation and moral assessment [8, 15, 14, 20, 17, 18]) have assumed that reputations are objective and common knowledge in the population – meaning that opinions about reputations do not differ among individuals. Here we relax this assumption and allow individuals to differ in their opinions about one another. This reveals an under-appreciated subtlety in the application of norms for updating reputations. Namely, when an observer updates the reputation of a donor interacting with a recipient, the “reputation of the recipient” could be considered either from the observer’s own perspective, or from the donor’s perspective. Under a purely egotistical application of a social norm, the “recipient reputation” means the reputation in the eyes of the observer, who is forming an assessment of the donor. In this case the observer either ignores, or is unaware of, the donor’s view of the recipient. This case corresponds to *E* = 0 in our analysis, the no-empathy model of moral assessment. However, we also analyze the possibility of empathetic moral assessment, *E* > 0, whereby the observer may account for donor’s view of the recipient’s reputation when assessing the donor. In the extreme case *E* = 1, for example, the observer always uses the donor’s view of the recipient’s reputation when applying the social norm to update the donor’s reputation (Figure 1).

### 2.2 Empathetic moral judgment induces cooperation

To analyze how empathy influences cooperation we first examine strategy evolution with a fixed degree of empathy 0 ≤ E ≤ 1. We use the classic replicator-dynamic equations [21, 9] that describe how the frequencies of strategies (ALLD, ALLC, and DISC) evolve over time in an infinite population of players’ strategies reproducing according to their payoffs. For each of the four most common norms we find bi-stable dynamics [20]. That is, depending on the initial conditions the population will evolve to one of two stable equilibria: a monomorphic population a pure defectors, which supports no cooperation, or a population of cooperative (non-ALLD) strategies that supports some cooperation.

How does empathy influence the prospects for cooperation? Under the Scoring norm, strategic evolution does not depend on the degree of empathy, because this norm ignores the recipient’s reputation when updating a donor’s reputation. For all the other norms considered, however, empathy tends to increase cooperation. In particular, the basin of attraction towards the stable equilibrium that supports cooperation (green regions in Figure 2) is always larger when players are more empathetic – meaning that when *E* is larger, there is a larger volume of initial conditions in the strategy space that lead to the stable equilibrium supporting cooperation.

**Figure 2:**
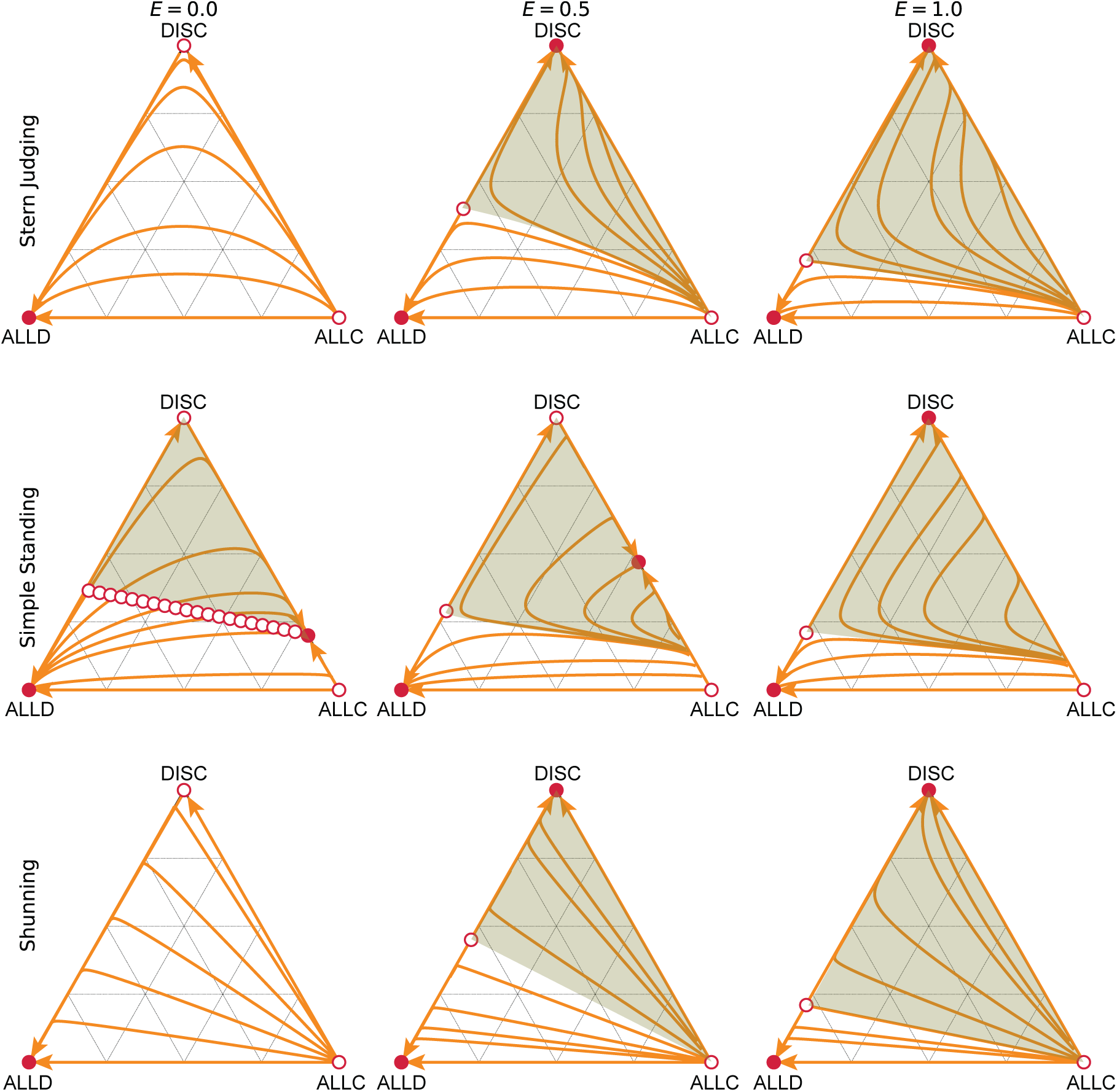
Empathetic moral evaluation facilitates the evolution of cooperation. We analyzed strategy evolution in the donation game under different social norms of moral assessment. Triangles describe the frequencies of three alternative strategies: unconditional defectors (ALLD), unconditional cooperators (ALLC), and discriminators (DISC) who cooperate with good recipients and defect against bad recipients. Red circles indicate the stable (filled) and unstable (open) strategic equilibria under replicator dynamics. The basin of attraction towards a stable equilibrium that supports cooperation (green) is larger as empathy, E, increases, for all three social norms shown. Orange curves illustrate sample trajectories towards stable equilibria. Costs and benefits are *c* = 1.0, *b* = 5.0, and error rates are *e*_1_ = *e*_2_ = 0.02.

In the case of Shunning and Stern judging, the stable equilibrium that supports cooperation consists of a monomorphic population of discriminators (Figure 2). Not only is the basin of attraction towards this equilibrium larger when a population is more empathetic, but so too is the equilibrium frequency of cooperative actions increased by greater degrees of empathy (Figure 2 and Supplementary Figure 1). And so empathy increases the frequency of outcomes that support cooperation, and also increases the frequency of cooperation at these outcomes.

In the case of Simple Standing, the stable equilibrium that supports cooperation consists of a mix of ALLC and DISC strategists. The discriminator frequency at this equilibrium increases with empathy as

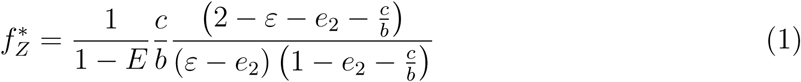

where *ε* = (1 — *e*_1_)(1 — *e*_2_) + *e*_1_*e*_2_, until it reaches 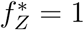. The rate of cooperative play at this mixed equilibrium shows only a weak dependence on the degree of empathy (Supplementary Figure 1).

Aside from the stable equilibria discussed above, for all four norms there is also an unstable equilibrium, with some portion of the population playing ALLD and some portion playing DISC. The frequency of discriminators at this unstable equilibrium is

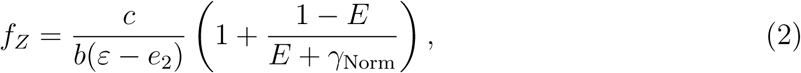

where 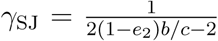,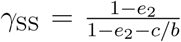and 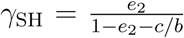,*γ*SC → ∞. When *E* = 1 these results coincide with the expressions found by Sasaki et al. [20]. This result reflects the sense in which previous studies, assuming no variation in personal opinions about reputations, are mathematically equivalent to always taking another person’s perspective (*E* = 1).

### 2.3 Social norms that promote cooperation

In a finite population the frequencies of strategies do not evolve towards a fixed stable equilibrium, but rather continue to fluctuate once they reach stationary state, irrespective of initial conditions, due to demographic stochasticity. To study the impact of empathy on cooperation in this setting we undertook Monte Carlo simulations. In this model, successful strategies spread through social contagion: a strategy is copied with the probability 1/(1+exp(−*w*[∏_1_ − ∏_0_])), where *w* is the selection strength, and ∏_1_ and ∏_0_ are payoffs of two randomly selected individuals ([22, 23], see Methods). In addition to these imitation dynamics, player strategies also change via random exploration at a rate *μ*. For the sake of simplicity, we assumed that the timescale at which games are played and payoffs are acquired is much faster than the timescales of imitation, exploration, and reputation dynamics, so that each individual plays many games before any of these events take place (see Methods).

Empathy tends to increase mean levels of cooperation in finite populations under stochastic dynamics (Figure 3), similar to our findings in an infinite population. The effects of empathy are pronounced: the stationary mean frequency of cooperation ranges from near zero to near unity, in response to increasing the value of the empathy parameter *E*.

**Figure 3:**
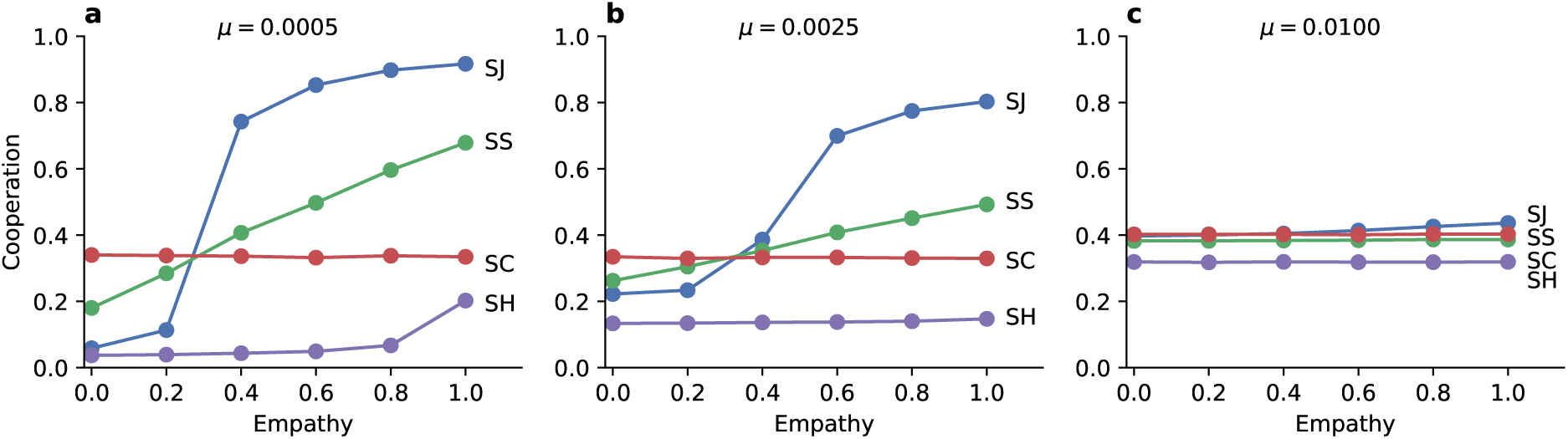
Empathetic moral judgment increases the average rate of cooperation under simple social norms. The degree of empathy, *E*, determines which social norms of moral assessment produce the most cooperation and thus greatest social benefit. (**a**) The Stern Judging (SJ) norm supports the highest rates of cooperation when the value of empathy is high. But Stern Judging performs poorly under egocentric moral judgment, where Scoring (SC) and Simple Standing (SS) dominate. The scoring norm (SC) does not depend on reputation, and so it shows cooperation levels that are insensitive to empathy. The Shunning norm (SH) always produces the lowest level of cooperation. (**b, c**) As the strategy exploration rate *μ* increases, SJ and SS become less efficient at promoting cooperation under highly empathetic moral evaluations, but preform better under egocentric judgment of observations. The figure shows ensemble mean cooperation levels in replicate Monte Carlo simulations of *N* = 100 individuals.

For high values of empathy, Stern Judging is the most efficient social norm at promoting cooperation, followed by Simple Standing and Scoring. This rank ordering of social norms is consistent with the prior literature [17, 18, 19]. However we find a striking reversal from the established view of social norms when individuals are less empathetic. As *E* → 0 Scoring promotes the most cooperation, while Stern Judging and Simple Standing engender less cooperation. And so the level of empathy strongly influences the amount of cooperation that evolves, and it even changes the ordering of which social norms are best at promoting cooperation. In particular, Stern judging is the most socially beneficial norm [15, 17, 18, 19] only when individuals account for subjectivity in moral assessment, or when individuals are forced to agree with one another through a centralized institution of moral assessment.

### 2.4 Evolution of empathy

We have seen that empathy promotes cooperation in finite populations with reputation-dependent strategies. It remains unclear, however, if empathy itself can evolve to high levels, and whether a population of empathetic individuals can resist invasion from egocentric moral evaluators. In the following analysis we assume that the degree of empathy in moral evaluation can be observed, inferred or learned, and can therefore evolve through social contagion (imitation dynamics). Alternatively, an individual’s capacity for empathetic observations may have a genetic component evolving via Darwinian selection [24].

We analyze the evolution of empathy using the framework of adaptive dynamics [4]. Assuming rare mutations to the continuous empathy trait *E* ∈ [0,1], we calculate the invasion fitness of an invader *E*_1_ in an infinite resident population with empathy *E*_R_ by comparing their expected payoffs. We report pairwise-invisibility plots and investigate the evolutionary stability of singular points *E*\*, where the gradient of invasion fitness *∂W*(*E*_R_, *E*_1_)/*∂E*_1_ vanishes. To support our analytic treatment we also perform Monte Carlo simulations in finite populations subject to demographic stochasticity, where empathy evolves through social copying according to individual payoffs, similar to strategy evolution under imitation dynamics [22].

Evolution can often favor empathy, depending upon the social norm and the initial conditions. To study empathy dynamics, we initially assume that the population is monomorphic for the discriminator strategy. In the case of the Shunning norm, then, there is a single, repulsive singular value of empathy (Figure 4e) at

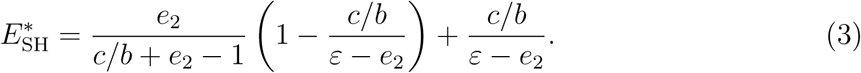

**Figure 4:**
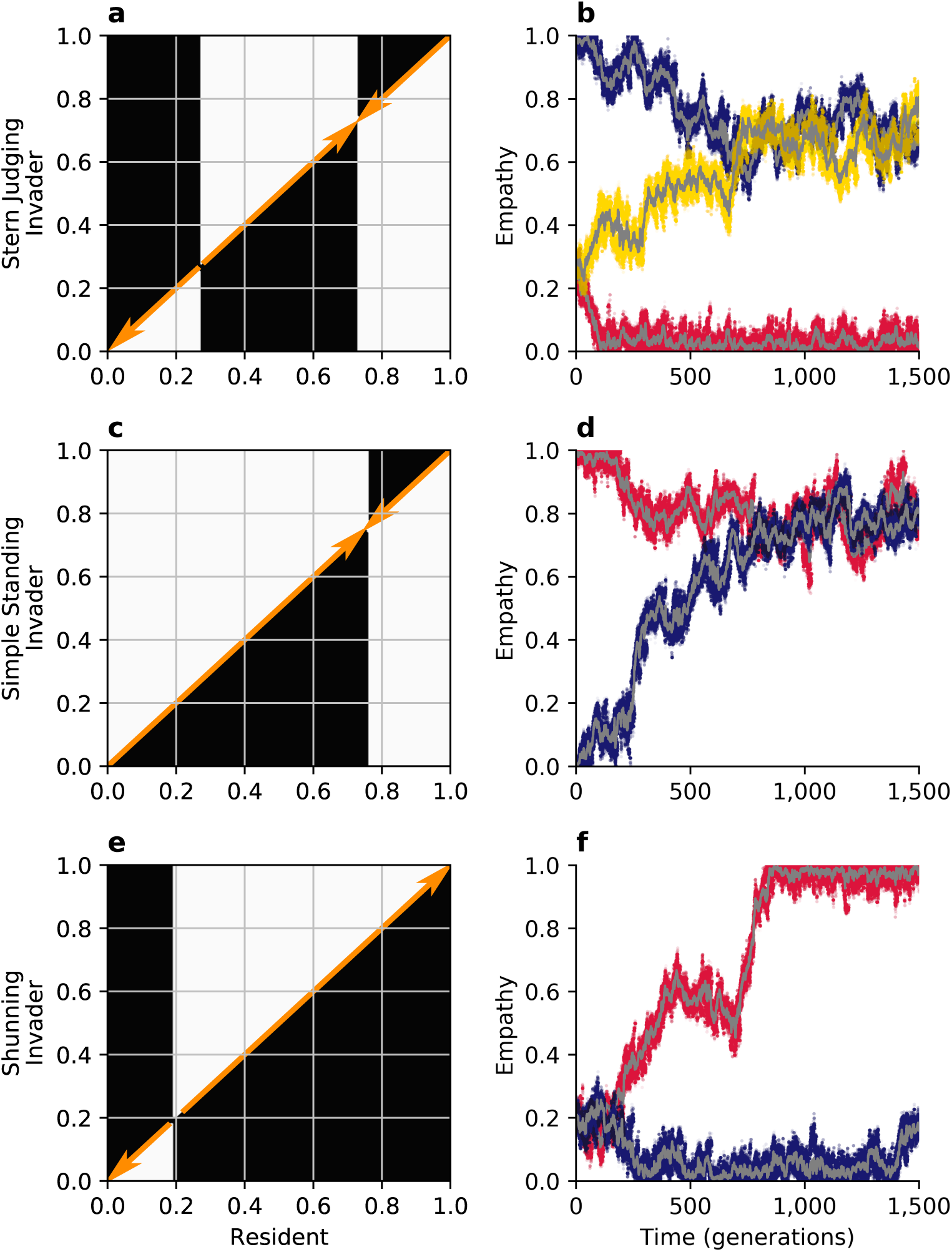
Evolution of empathy, *E*, in a population of discriminators under three different social norms. (**a, c, e**) White areas in the pairwise invasibility plots show values of *E* for which the invader’s expected payoff exceeds the mean payoff of the resident population. Orange arrows indicate the direction of evolution in an infinite population. (**b, d, f**) Monte Carlo simulations in small populations of 100 individuals with recurring mutations to *E* reflect the predictions of adaptive-dynamics analysis. Sample trajectories showing all *E* values in three different populations are show in colors (red, blue, yellow), with their population means shown in gray.

Such a population is bistable. If the initial level of empathy exceeds 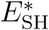 the population will evolve towards complete empathy (*E* = 1) and the discriminator strategy will remain stable; but if the initial level of empathy is less than 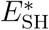 the population will evolve towards complete egocentrism (*E* = 0), at which point the discriminator strategy is no longer stable (Fig 2g) and the population will be replaced by pure defectors. The singular value 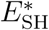 decreases as the benefit of cooperation *b*/*c* increases, permitting a larger space of initial conditions that lead to the evolution of complete empathy (Figure 5c). And so, in summary, under the Shunning norm long-term strategy and empathy co-evolution will tend towards a completely empathetic population of discriminators, especially when the benefits to cooperation are high; or, alternatively, evolution will lead to complete population-wide defection.

**Figure 5:**
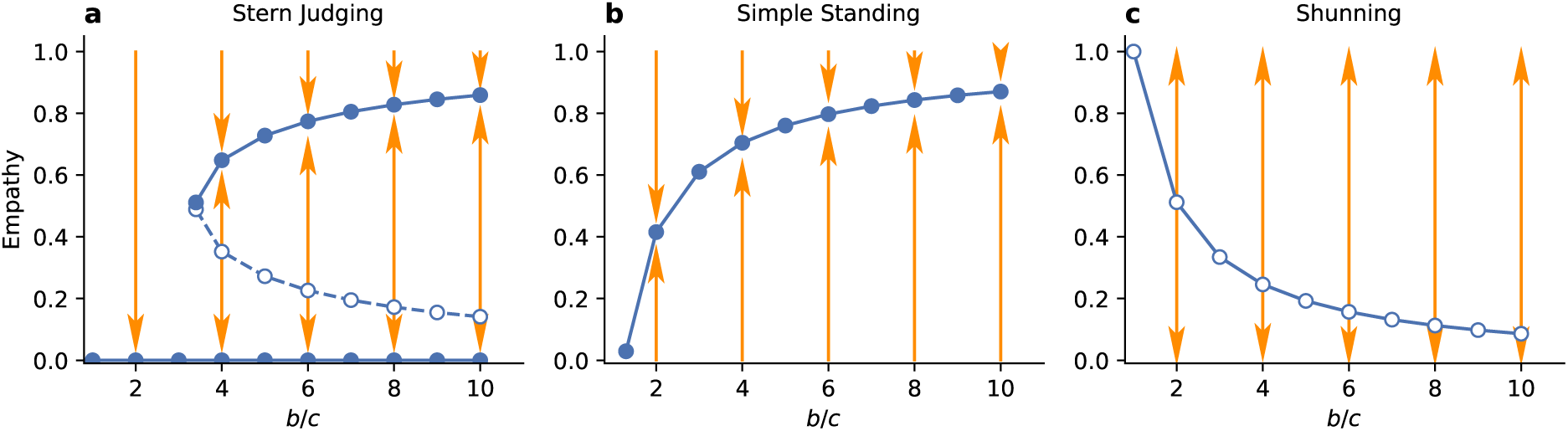
Evolution of empathy in an infinite population of discriminator strategists. Circles indicate evolutionarily stable (solid) and unstable (open) singular values of empathy, *E*. (**a**) Above a critical benefit-cost ratio *b*/*c*, increasing the benefit of cooperation promotes the evolution of high levels of empathy under Stern Judging norm. (**b**) Under Simple Standing there is a single ESS value for empathy. The highest levels of empathy evolve with high benefits and low costs of cooperation. However, in this case the monomorphic discriminator equilibrium is not stable at the ESS value of empathy. (**c**) In populations governed by the Shunning norm there are no stable internal equilibria for *E*, and empathy will evolve to either one or zero.

Similar dynamics occur under the Stern Judging norm. In this case, starting from a monomorphic population of discriminators, there are two singular values for *E*: an evolutionary repeller 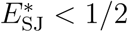 and attractor 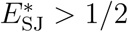 (Figure 4a, b) given by

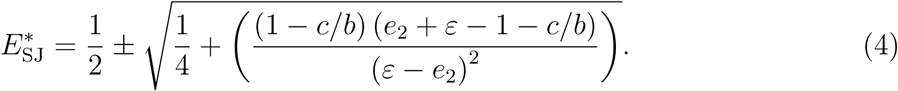

Provided empathy initially exceeds the repulsive value evolution will favor increasing empathy towards the attractive value, and the population of discriminators will remain stable. Increasing *b*/*c* again favors the evolution of empathy, as it increases the value of the locally stable 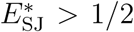 and also the range of initial values that that lead to 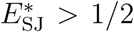 through fixation of small mutations (Figure 5a). However, if empathy starts below the repulsive value, selection will favor evolution toward the attractive singular point at 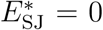, which no longer supports DISC as a stable equilibrium in strategy space (Figure 2a). And so, in summary, under Stern Judging co-evolution of strategies and empathy will tend towards a highly empathetic population of discriminators, especially when the benefits to cooperation are large; or, alternatively, evolution will lead to all defectors and empathy will thereafter drift neutrally.

The evolution of empathy is more complicated under the Simple Standing norm. Assuming the population consists of discriminators there is a single evolutionarily stable and globally attractive singular point 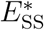 (Figure 4c). The value of empathy at this singular point is once again larger when the benefits of cooperation are larger (Figure 5b). However once this value of empathy is reached, the strategic equilibrium at pure discriminators is no longer stable, and the population will instead be replaced by a mix of DISC and ALLC strategists, under replicator dynamics. This new strategic equilibrium will, in turn, lower the singular value of 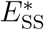 under adaptive dynamics, which again changes the equilibrium balance of DISC and ALLC strategists. Long-term strategy-empathy co-evolution will continue in this fashion, with both ALLC and DISC present the population, until the singular value of empathy reaches 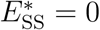 (Supplementary Figure 2). The strategic equilibrium at this point lies on the boundary of two basins of attraction (Figure 2d) and is vulnerable to invasion by pure defectors in a finite population. And so, in summary, while the exact dynamics will depend on the time scales of empathy and strategy evolution, Simple Standing cannot sustain empathy over the long term as these both components of personality evolve, eventually resulting in a population of pure defectors.

## 3 Discussion

Empathy has long been associated with altruism in humans. Existing theory focuses on linkage of emotional states between individuals and empathy-induced helping [1, 3, 7]. For instance, there is substantial evidence that the capacity to take another person’s perspective increases empathetic concern and promotes ‘truly selfless’ altruism [1]. By contrast, here we treat the evolution of empathetic perspective-taking in the context of moral evaluation. Understanding the impact of empathy on cooperation in this context is critical to understanding its role in modern highly-connected human societies that lack a central institution of objective moral assessment.

Social norms specify the rules of moral evaluation. It is well known that moral reputations can sustain high levels of cooperation if individuals discriminate between the ‘good’ and the ‘bad’. Social norms themselves likely emerge from individual beliefs of what reputations should be assigned to defectors and cooperators in distinct social situations. While some argue that social reputations are absolute – for instance due to shared information, institutions and gossip – our study draws attention to the potential for disagreements on reputations that arise from errors or different observation histories. The same individual can have different reputations in the eyes of distinct observers; in other words, moral evaluations are not absolute, and social reputation is relative.

One of our key findings is that high levels of cooperation can be sustained only if individuals recognize this moral relativity and are capable of making moral judgments from another person’s perspective. Egocentric world-views lead to unjustified or irrational defection, because a person perceived as ‘bad’ by the observer might actually appear ‘good’ in the eyes of the donor who’s action is being evaluated, or *vice versa*. This point is particularly striking in the case of Stern Judging, the norm that assigns a ‘good’ reputation only to individuals who cooperate with other ‘good’ players and defect against ‘bad’ [6, 15]. Despite being the most efficient norm at promoting cooperation in empathetic societies, Stern Judging performs very poorly in egocentric populations. On the other hand, Scoring – the norm that does not take into account the recipient’s reputation at all – is immune to the effects of empathy and dominates in societies with egocentric moral evaluation rules.

Finally, we have shown that high levels of empathy in moral evaluation can evolve through cultural copying, and remain evolutionarily stable if the society is governed by Stern Judging or Shunning norms. Once these societies evolve empathy, individuals performing egocentric evaluations of observed social behavior will be rewarded less than their empathetic peers, and this remains true even if strategies are allowed to co-evolve with empathy.

However, we have also shown that under all four social norms, egocentric and uncooperative societies are nevertheless possible evolutionary outcomes. In populations governed by Stern Judging, Shunning and Scoring this outcome represents an alternative locally (though not globally) attractive stable state in the strategy-empathy phase space. In the case of Simple Standing, by contrast, the egocentric and uncooperative outcome is the only long-term stable outcome as both empathy and strategies are allowed to evolve.

Our study of the role of empathy in human cooperation raises a number of questions to be addressed in future work. One question involves the competition of social norms for moral evaluation – a topic that has been studied in a few restrictive contexts, such as when errors do not occur [25], or in the context of population structure [15]. Perhaps an even more important question is whether and how population-wide social norms can evolve from individual moral beliefs to begin with. It is unclear whether social contagion or individual-level Darwinian selection is sufficient to establish a hierarchy of norms governing individual behavior in a population. We have shown that the norms that promote the most cooperation change depending on the individual capacity for empathetic perspective-taking, but should we also expect different norms to evolve under empathetic and egocentric modes of judgment? For instance, populations characterized by fully empathetic moral judgment might be conducive to the evolution of selfish norms that indiscriminately assign ‘bad’ reputations to evade costly cooperation without being punished, while models with private egocentric evaluation may lead to the evolution of more cooperative norms, such as Scoring or Stern Judging ([26, 25]).

## 4 Methods

### 4.1 Cooperation under empathetic moral evaluation

#### 4.1.1 Replicator dynamics

To analyze evolutionary dynamics in the strategy space, we consider replicator dynamics in infinite populations with fixed social norms and fixed values of the empathy parameter *E*, limiting ourselves to the discrete three-strategy space (ALLC or *X*, ALLD or *Y* and DISC or *Z*). Denoting the mean payoff of strategy *s* as ∏_*s*_, and the frequencies of the three strategies at *f_s_*, the strategy evolution dynamics can be described as 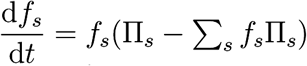.

To describe image dynamics, we let *g* denote the frequency of ‘good’ individuals within the population, i.e. *g* = *fXgX* + *fYgY* + *fzgZ*. For the Stern Judging norm, we have

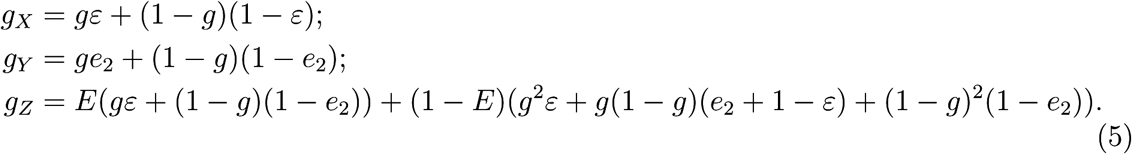

Here *ε* = (1 — *e*_1_)(1 — *e*_2_) + *e*_1_*e*_2_. Expected payoffs of the three strategies are then:

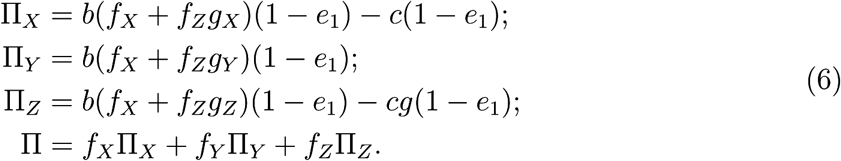

Likewise for Shunning:

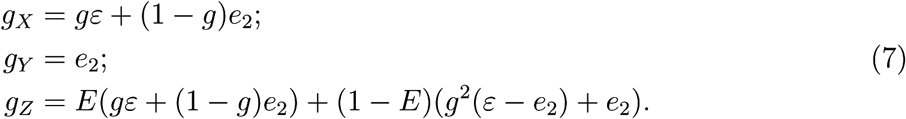

For Simple Standing:

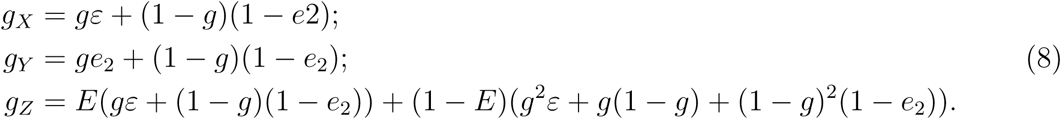

And finally for Scoring norm, empathy *E* is irrelevant, because the norm does not take into account the reputation of the recipient:

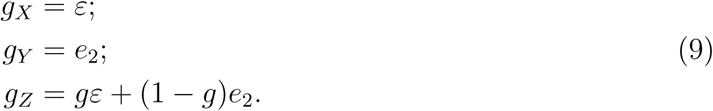

#### 4.1.2 Stochastic simulations

In addition to the deterministic replicator-dynamics analysis of strategy evolution, we performed a series of individual-based simulations to measure mean levels of cooperation under continuous influx of mutations in the strategy space ([17]). We assume that all individuals follow the same social norm and are characterized by the same value of empathy, *E*. The population consists of *N* individuals, each with its own strategy and its own subjective list of reputations. Each generation, any given individual interacts with all other members of the society in three different roles: once as a donor, once as a recipient, and once as an observer.

First, each individual plays a single round of the donation game with all other members of the society according to her strategy *S* = [*p*, *q*] and the subjective reputation of the recipient, also taking into account the implementation error *e*_1_. Here *p* and *q* denote the probabilities that a donor cooperates with a ‘bad’ (B) and ‘good’ (G) recipient, respectively. The act of cooperation fails with the probability *e*_1_ (defection always succeeds). The cumulative payoff is then assigned to each individual, with the benefit of cooperation fixed at *b* and the cost of a cooperative act *c*.

To update their list of subjective reputations based on the social norm *N_ij_*, each player then chooses to observe a single interaction per donor (that is, with a randomly chosen recipient), again taking into account subjective reputation of the recipient either in the eyes of the donor (probability *E*) or the eyes of the observer (with a probability 1 — *E*). The newly assigned reputation is reversed with the probability *e*_2_, representing observation errors. For the sake of simplicity, we assume that all reputations are updated simultaneously after all donor-recipient interactions have taken place.

We model selection and drift of strategies as a process of social contagion implemented as a pairwise comparison process. Following the reputation-updating step, a random pair of individuals is chosen; the first individual adopts the strategy of the second with the probability 1/ (1 + exp(—*w*[∏_1_ – ∏_0_])), where *w* is the selection strength, and ∏_1_ and ∏_0_ are payoffs of the two earned within the last generation. In our simulations of populations with *N* = 100 individuals, we used *w* = 1.0. Finally, each individual is subject to random strategy exploration, in which a new random strategy is adopted with a small probability *μ* ([17])

The simulation is initialized with random strategies and random lists of subjective reputations. We recorded the mean rate of cooperation averaged over 150,000 generations in 50 replicate populations, which is reported in Figure 3.

### 4.2 Evolution of empathy

Let *g_ij_* be the frequency of ‘good’ individuals in the sub-population *i* as seen by individuals belonging to the sub-population *j*, where *i* and *j* correspond either to resident (*i*, *j* = 0) or invader (*i*, *j* = 1) sub-population. Working in the limit of negligible invader frequencies, for Stern Judging norm we have:

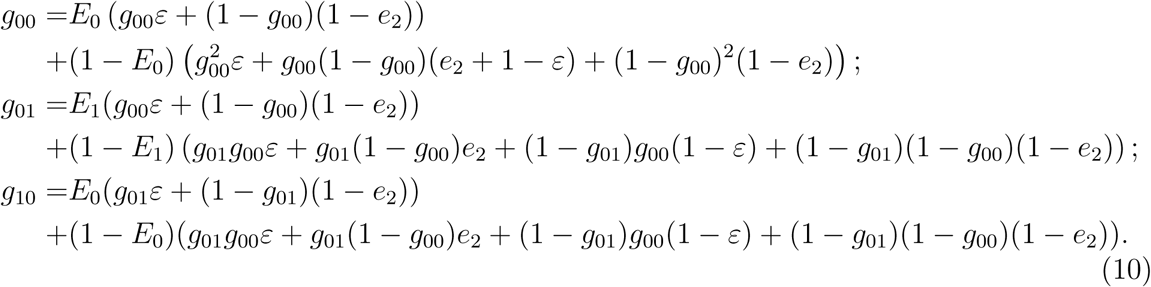

Here *ε* = (1 — *e*_1_)(1 — *e*_2_) + *e*_1_*e*_2_, and *E*_0_ and *E*_1_ are empathy values of resident and invader sub-population. For Simple Standing norm, the relative frequencies of good individuals are:

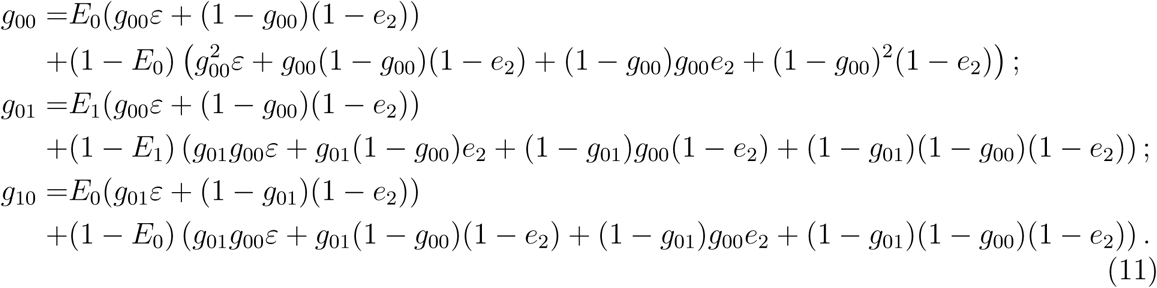

Likewise, for the Shunning norm:

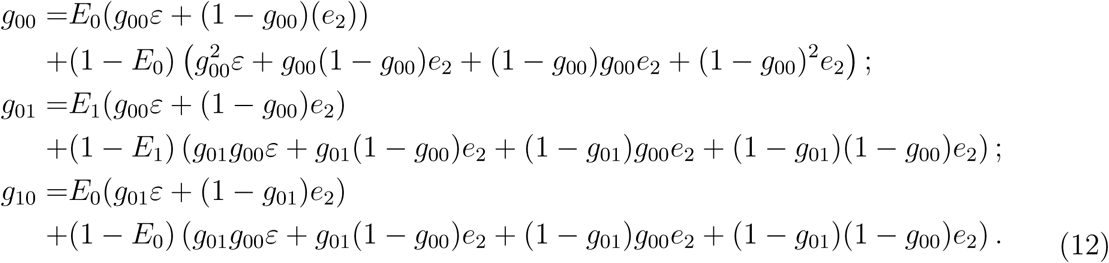

Under Scoring, the frequencies of ‘good’ individuals do not depend on empathy:

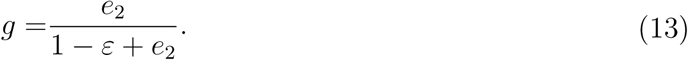

We then calculate the expected payoffs of individuals in resident and invader sub-populations:

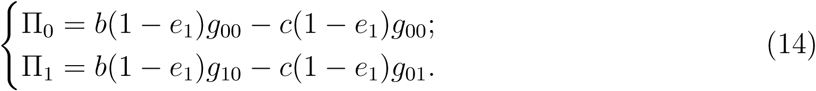

These payoffs are used to generate pairwise invasibility plots in Figure 4. Singular points are found by setting 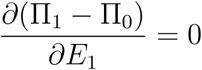 and setting *E*_0_ = *E*_1_

### 4.3 Individual based simulations of empathy evolution

To verify the ESS results of the adaptive-dynamics calculations we performed a series of Monte-Carlo simulations in finite populations of *N* = 100 individuals. The simulation routine is largely the same as for the strategy evolution (section 4.1.2), except that in this case we fixed the strategy at DISC and allowed *E* to evolve via constant influx of small mutations. Each generation, empathy of an individual changes via mutation at a rate *μ_E_* = 0.005. Since empathy is a continuous parameter, we draw the mutational deviation *δ_E_* from a normal distribution centered around *δE*_0_ = 0 with a standard deviation *σ* = 0.01. Selection for empathy is modeled by choosing 5 random pairs of individuals and assuming that in each pair the first individual copies the empathy value *E*_1_ of the second with the probability 1/ (1 + exp(–*w*[∏_1_ – ∏_0_])), where ∏_1_ and ∏_0_ are their payoffs.

## 5 Supplementary Figures

**Supplementary Figure 1:**
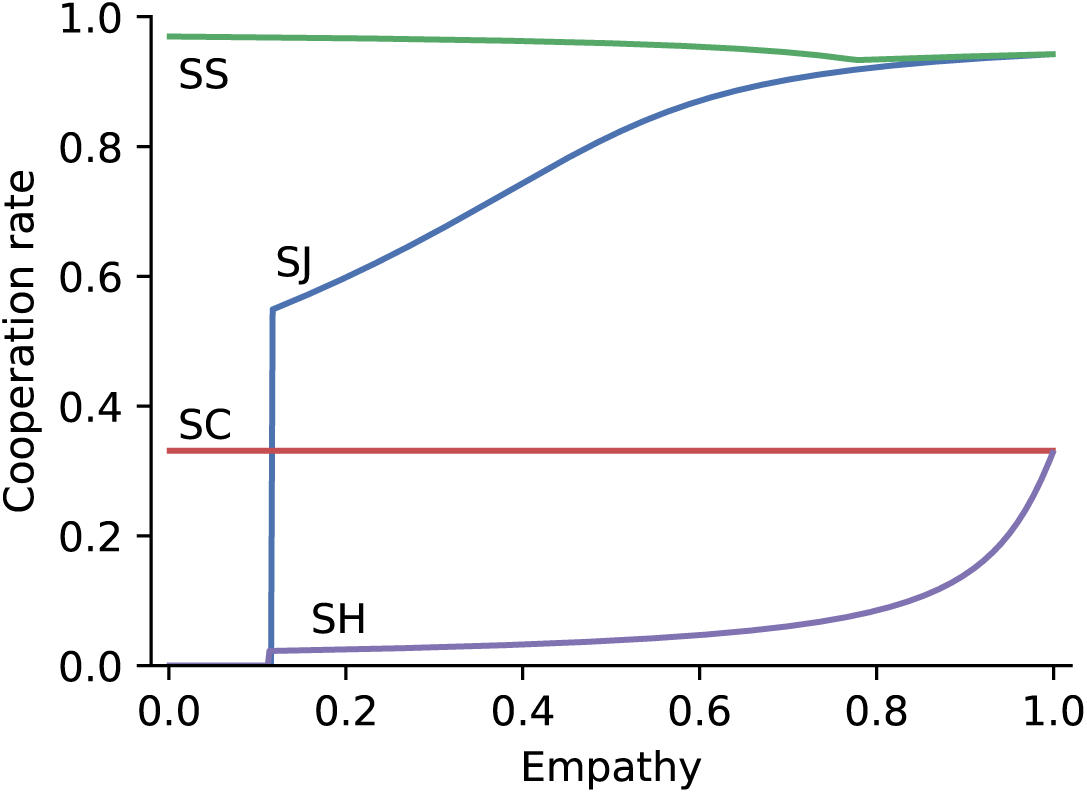
Cooperation rates at the cooperative equilibria in the replicator dynamics of strategy evolution. For Stern Judging (SJ), Scoring (SC) and Shunning (SH) norms the cooperative equilibrium (where it exists) corresponds to a homogeneous population of DISC players, while for Simple Standing (SS) it consists of a mixture of ALLC and DISC strategists. *b*/*c* = 5, *e*_1_ = *e*_2_ = 0.02.

**Supplementary Figure 2:**
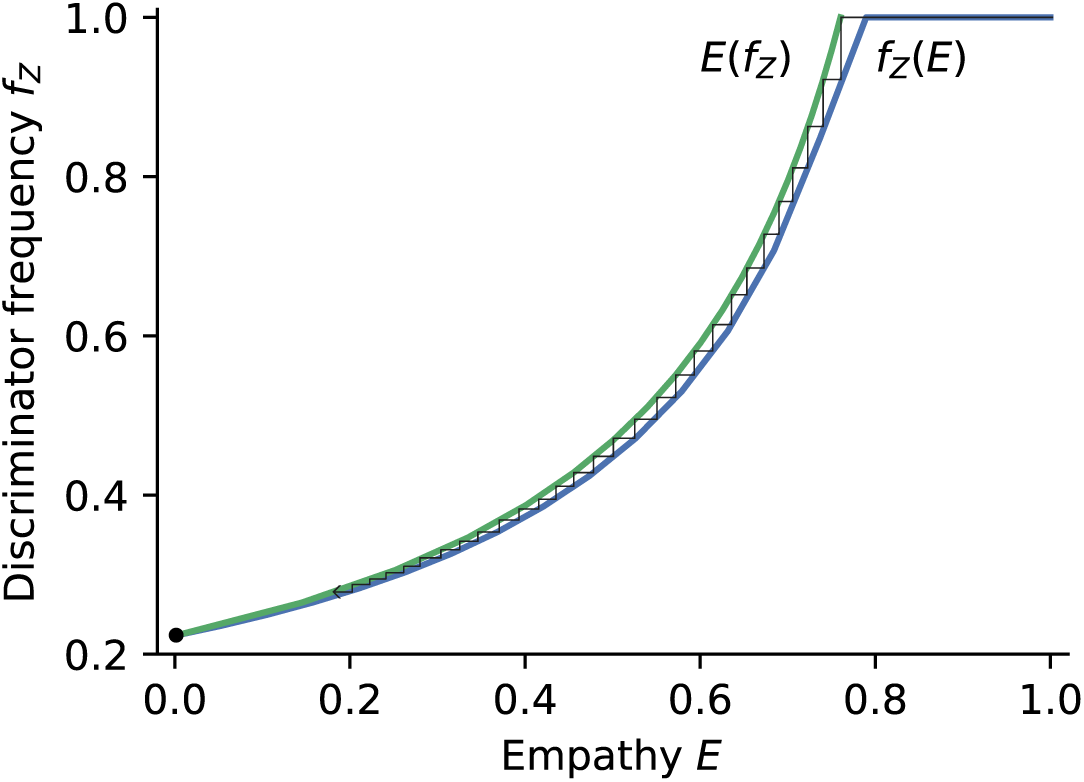
Empathy-strategy co-evolution in an infinite population under Simple Standing norm. The figure combines the equilibria analysis in the replicator dynamics for strategy evolution given fixed empathy, *f_Z_*(*E*); and singular-point analysis in pairwise invasibility plots for empathy evolution, *E*(*f_Z_*). We make no assumption about the timescales of the two mutational processes. The only fixed point across the two models is *E* = 0.

